# Kaposi’s sarcoma-associated herpesvirus (KSHV) gB dictates a low-pH endocytotic entry pathway as revealed by a dual-fluorescent virus system and a rhesus monkey rhadinovirus expressing KSHV gB

**DOI:** 10.1101/2023.11.07.566061

**Authors:** Sarah Schlagowski, Anna K. Großkopf, Shanchuan Liu, Natalia Khizanishvili, Stefano Scribano, Armin Ensser, Alexander S. Hahn

## Abstract

Interaction with host cell receptors initiates internalization of Kaposi’s sarcoma-associated herpesvirus (KSHV) particles via endocytosis or macropinocytosis. Fusion of viral and host cell membranes, which is followed by release of the viral capsid into the cytoplasm, is executed by the core fusion machinery composed of gH, gL, and gB, that is common to all herpesviruses. KSHV infection has been shown to be sensitive to inhibitors of vacuolar acidification, suggestive of low pH as a fusion trigger.

To analyze KSHV entry at the single particle level we developed single- and dual-fluorescent recombinant KSHV strains that incorporate tagged glycoproteins, capsid proteins or a combination thereof. In addition, we generated a hybrid rhesus monkey rhadinovirus (RRV) that expresses KSHV gB in place of RRV gB to analyze gB-dependent differences in infection pathways.

Our data demonstrate lytic reactivation and infectivity of dual-fluorescent KSHV and incorporation of fluorescently-tagged proteins into viral particles. Confocal microscopy was used to quantify co-localization of fluorescently-tagged glycoproteins and capsid proteins. By using the ratio of dual-positive KSHV particles to single-positive capsids as an indicator of fusion events we established the KSHV fusion kinetics upon infection of different target cells.

We measured marked differences in the “time-to-fusion” between different cell types. Inhibition of vesicle acidification prevented virus-cell fusion, implicating low vesicle pH as a requirement for the KSHV fusion step. These findings were corroborated by comparison of RRV-YFP reporter virus with wildtype gB and RRV-YFP encoding KSHV gB in place of RRV gB. While RRV wt infection of receptor-overexpressing cells was unaffected by inhibition of vesicle acidification, RRV-YFP expressing KSHV gB was sensitive to Bafilomycin A1, an inhibitor of vacuolar acidification.

Single- and dual-fluorescent KSHV strains eliminate the need for virus-specific antibodies and enable the tracking of single viral particles during entry and fusion. Together with a hybrid RRV expressing KSHV gB, these two novel tools identify low vesicle pH as an endocytotic trigger for KSHV membrane fusion.

## INTRODUCTION

KSHV is a human oncogenic herpesvirus. It is associated with several malignancies: Kaposi’s sarcoma, multicentric Castleman’s disease, primary effusion lymphoma and potentially osteosarcoma ([1] reviewed in [2]). It is further associated with Kaposi Sarcoma inflammatory cytokine syndrome (KICS)[3,4].

Enveloped viruses including the herpesviruses employ two different strategies regarding the timing of membrane fusion: They either fuse directly at the cell surface with the plasma membrane or they fuse from the interior of endocytic vesicles with the membranes of these vesicles. For the first strategy, endocytosis is not necessary and potentially detrimental to infection if the environment in endocytic compartments is not conducive for fusion. For the second strategy, endocytosis is a prerequisite. As in most biological systems the distinction is usually not absolute and many viruses are capable of utilizing both strategies, but they usually fall predominantly into one or the other category. Often, the ability of viral glycoproteins to efficiently cause cell-cell fusion upon expression correlates well with the ability of the respective viruses to enter through fusion at the plasma membrane. An instructive example is SARS-CoV-2, which oscillates depending on the variant and its Spike protein between the two extremes. While the Delta variant Spike induces syncytia and mediates infection primarily through direct fusion, the BA.1 variant Spike does not induce syncytia and mediates entry primarily through endocytosis [5]. Among the herpesviruses, herpes simplex is a well-studied example for a virus with highly fusogenic glycoproteins that predominantly employs the direct fusion strategy but can also use the endocytic route in some circumstances [6–9]. KSHV on the other hand seems to fall in the second category [10–16].

KSHV uses a number of receptors for attachment and entry. It attaches to cells through binding to heparan sulfate proteoglycans like syndecans, which are engaged by glycoproteins K8.1, gB, KCP and gH [17–21]. gB additionally interacts with integrins [22,23], neuropilin 1 [11], and DC-SIGN [24,25]. Integrins and neuropilin-1, potentially also DC-SIGN, further promote endocytosis and/or macropinocytosis. KSHV also interacts with members of the Eph family of receptor tyrosine kinases, primarily with EphA2, through the gH/gL glycoprotein complex at a post-attachment step, and this interaction promotes endocytosis and/or macropinocytosis and fusion [10,12,15,26–32]. It has been observed that KSHV does not readily induce cell-cell fusion. While some groups managed to obtain a weak signal in cell-cell fusion assays [11,33,34], others did not [28,35]. Generally cell-cell fusion activity of KSHV glycoproteins is far below that of other viral glycoproteins, even from closely related viruses like the Epstein-Barr virus (EBV) [28] or the rhesus monkey rhadinovirus (RRV). These observations fit well with several publications that describe endo- or macropinocytotic entry of KSHV into target cells [10–16], and with reports describing inhibition of infection by substances that inhibit vesicle acidification [26,35,36]. An open question here is whether the inhibition of vesicle acidification has a direct effect on fusion of the viral and host cell membranes or whether it disrupts other processes like particle trafficking or post-entry signaling. We hypothesized that KSHV gB exhibits low fusogenicity at neutral pH and needs vesicle acidification for initiation of the fusion process. To test this hypothesis, we followed two approaches: i) we created a hybrid RRV and exchanged its more fusogenic gB with that of KSHV ii) we created a dual-fluorescent KSHV that allows to monitor the separation of viral envelope proteins and capsid as a surrogate marker for fusion.

## RESULTS

### Dual-fluorescent KSHV is replication-competent

We first confirmed the extremely low intrinsic fusogenicity of KSHV gB in cell-cell fusion assays, which is orders of magnitude below that of RRV (Fig. 1) and clearly necessitates a different approach to studying the fusion process.

**Figure 1.**
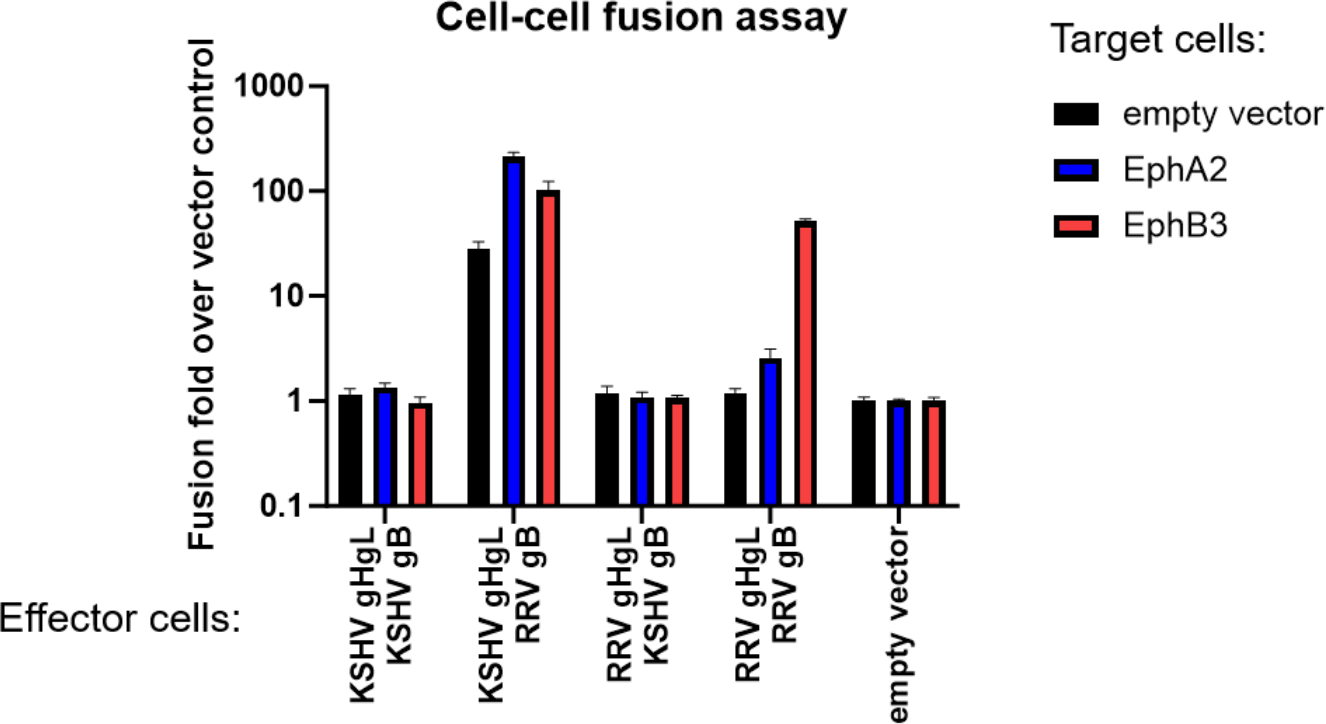
Low fusion activity of KSHV gB. Raji target cells were transduced with expression constructs for the indicated receptor proteins and were co-cultured with HEK293T effector cells transfected with expression constructs for the indicated viral glycoproteins or an empty vector. Fusion was quantified by luciferase readout. Error bars represent the standard deviation, the experiment was performed in triplicates.

We previously reported a KSHV reporter virus with a N-terminal mNeonGreen added to the capsid protein orf65, which had resulted in replication-competent virus [35], which can be tracked on a single particle level without the need for virus-specific antibodies. Based on this approach we constructed virus mutants harboring either an N-terminal mScarletH-tagged orf65 alone or in combination with mNeonGreen at the C-terminus of orf39, which encodes glycoprotein M (gM) (Fig. 2 A).

**Figure 2.**
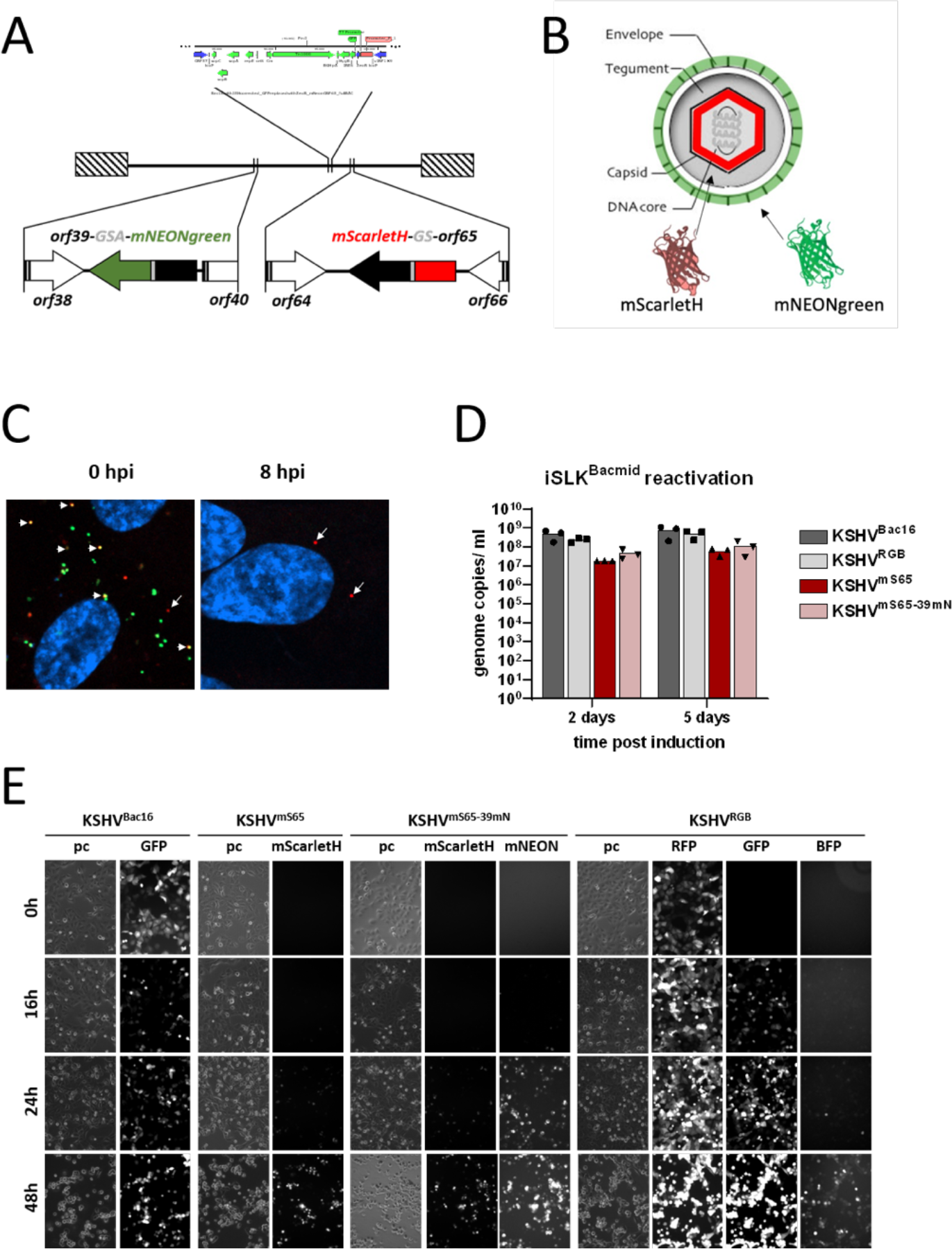
Construction of dual-fluorescent KSHV. A) Map of BAC 16 noGFP _mScarletH-ORF65_ORF39-mNEONgreen encoding KSHV^mS65-39mN^. B) Schematic drawing to illustrate the composition of dual-labeled KSHV particles. C) HUVEC cells imaged after virus attachment 0 hours post infection (hpi) or at the 8 hpi timepoint. D) Release of encapsidated genomes after chemical induction of the lytic cycle in iSLK producer cells harboring the indicated viruses. E) Fluorescence microscopy of iSLK cells that were infected with the first passage of the indicated viruses, selected and subjected to chemical induction of the lytic cycle. The cells were imaged in culture without fixation at the indicated timepoints after induction of the lytic cycle.

This led to double fluorescent viruses, labelled at the envelope through gM and at the capsid through ORF65 (Fig. 2 B). As expected, when imaged directly after addition to HUVEC cells, most red particles representing viral capsids were colocalized with green signal representing viral envelope. Of note we did detect a large number of single-mNEON+ spots without associated mScarletH signal. By 8h post infection mostly red capsids were visible, devoid of the green virus envelope. When we compared reactivation and release from producer cells of these tagged viruses to that of BAC16 or KSHV BAC16 RGB based viruses [37], release of viral particles was reduced by addition of mScarletH to ORF65. The reduction in virus yield was in the order of 1 log10. Introducing mNEONgreen fused to gM did not further impact virus yield (Fig. 2 D). The virus particles were infectious and fully replication competent as proven by another passage of the released virus and subsequent reactivation (Fig. 2 E). As the idea was to study fusion events and not replication over time, we decided to accept a reduction in virus yield, which probably is to be expected when adding several hundred copies, according to the literature between 800 and 960 copies [38], of a fluorescent protein to the virus capsid. We proceeded to use these viruses to study the fusion of KSHV with cellular membranes during entry into target cells.

### Fusion kinetics differ between different cells

We first analyzed the KSHV fusion kinetics in different types of cells. We performed time-lapse experiments to assess the time it would take for the envelope signal to dissociate from the capsid, a surrogate marker for virus-cell fusion. We chose SLK cells, an epithelial cell line, human umbilical vein endothelial cells (HUVEC), and human foreskin fibroblasts (HFF). We found that KSHV exhibited different fusion kinetics when infecting the different cells. Fusion occurred fastest in HUVEC, followed by SLK cells, and considerably more slowly in HFF cells (Fig. 3 A).

**Figure 3.**
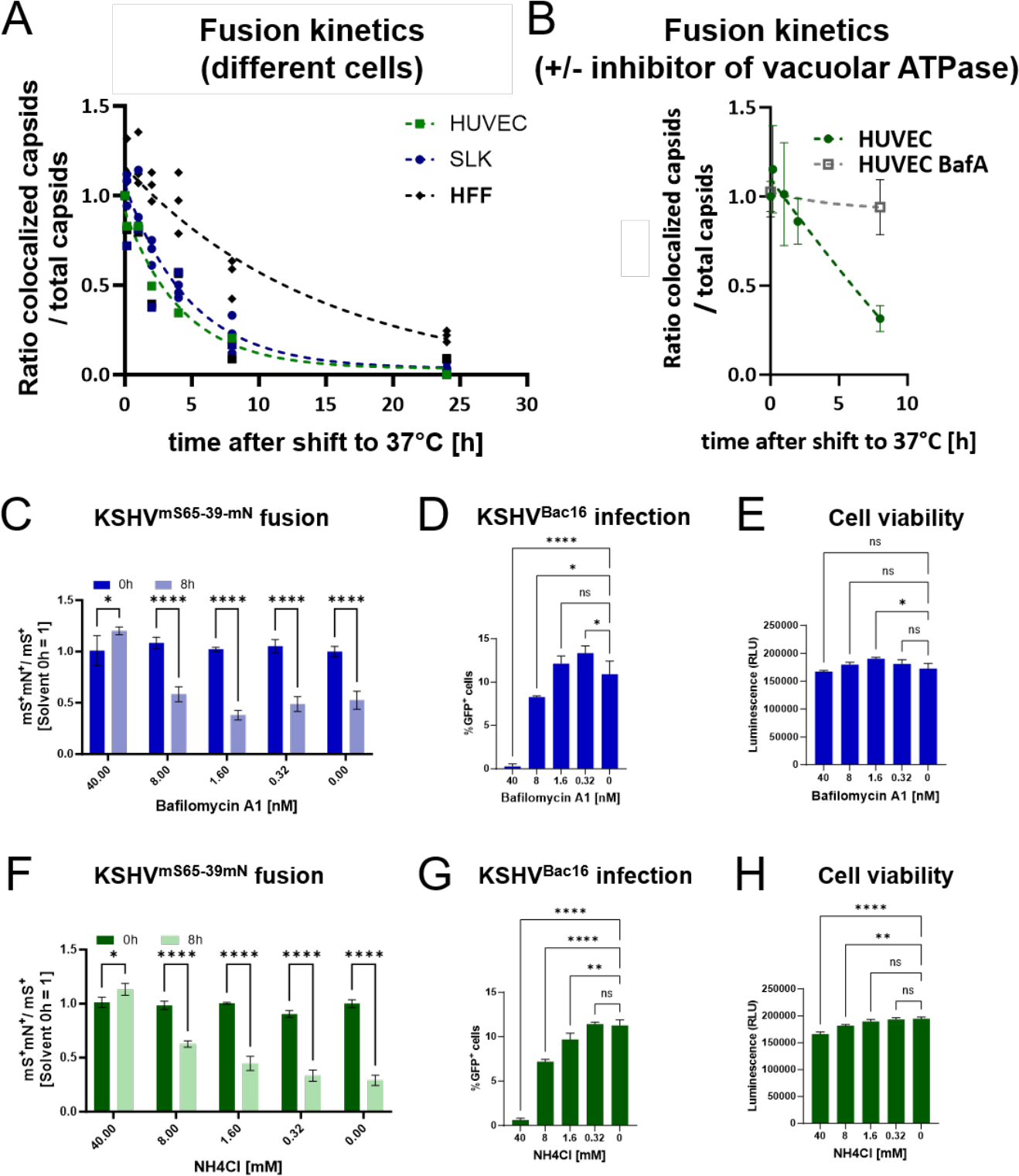
Cell-specific fusion kinetics and inhibition of fusion by inhibitors of vesicle acidification. A) Kinetics of virus-cell fusion with KSHV^mS65-39mN^ were plotted for HFF, HUVEC and SLK cells. The 0-hour-timepoint was set to 1 for all cells. The “Dissociation - One phase exponential decay” of GraphPad PRISM was used for non-linear curve fitting, error bars represent the standard deviation from three different images. Data represent the average of three independently produced virus stocks. B) Virus fusion with HUVEC was analyzed as in (A) but in the absence or presence of Bafilomycin A1 (BafA). C) Virus-cell fusion with SLK was analyzed as in (A) at the indicated timepoints and in the presence of the indicated concentrations of Bafilomycin A1. D) SLK cells were infected with KSHV BAC 16 in the presence of the indicated concentrations of Bafilomycin A1. E) SLK cells were incubated with Bafilomycin A1 and viability was measured. F) Virus-cell fusion with SLK was analyzed as in (C) in the presence of the indicated concentrations of ammonium chloride (NH4Cl). G) SLK cells were infected as in (D) in the presence of the indicated concentrations of ammonium chloride (NH4Cl). H) SLK cell viability was measured as in (E). Error bars represent the standard deviation, n=3. ns non-significant; * p<0.05; ** p<0.01; *** p<0.001, **** p<0.0001; two-way ANOVA with Sidak’s correction for multiple comparison (C, F), ordinary one-way ANOVA (D, E, G, H).

### KSHV virus-cell fusion is sensitive to inhibitors of vesicle acidification

We next tested whether fusion would be affected by inhibition of vesicle acidification. Our hypothesis was that the inhibitory activity of these substances against infection was based on their effect on virus-cell fusion. Indeed, we found that Bafilomycin A1 inhibited separation of envelope and capsids (Fig. 3 B & C), an inhibitor of vacuolar ATPase [39], and as expected also inhibited KSHV infection (Fig. 3 D) without toxicity (Fig. 3 E). Similar results were obtained with ammonium chloride (NH4Cl), a lysosomotropic agent that raises endosomal pH [40], which also prevented separation of capsid and envelope proteins (Fig. 3 F), inhibited infection (Fig. 3 G), but exhibited mild toxicity (Fig. 3 H).

We also controlled for the possibility that one of the two fluorescent proteins loses fluorescence more quickly under conditions of low pH, which would then result in an apparent loss of colocalization in case gM-mNeonGreen would fade faster than the mScarletH-orf65 in the capsid. When switching the position of the two fluorescent proteins, we observed the same separation of envelope proteins and capsid over time, and this again was inhibited by addition of Bafilomycin A1 (Fig. 4).

**Figure 4.**
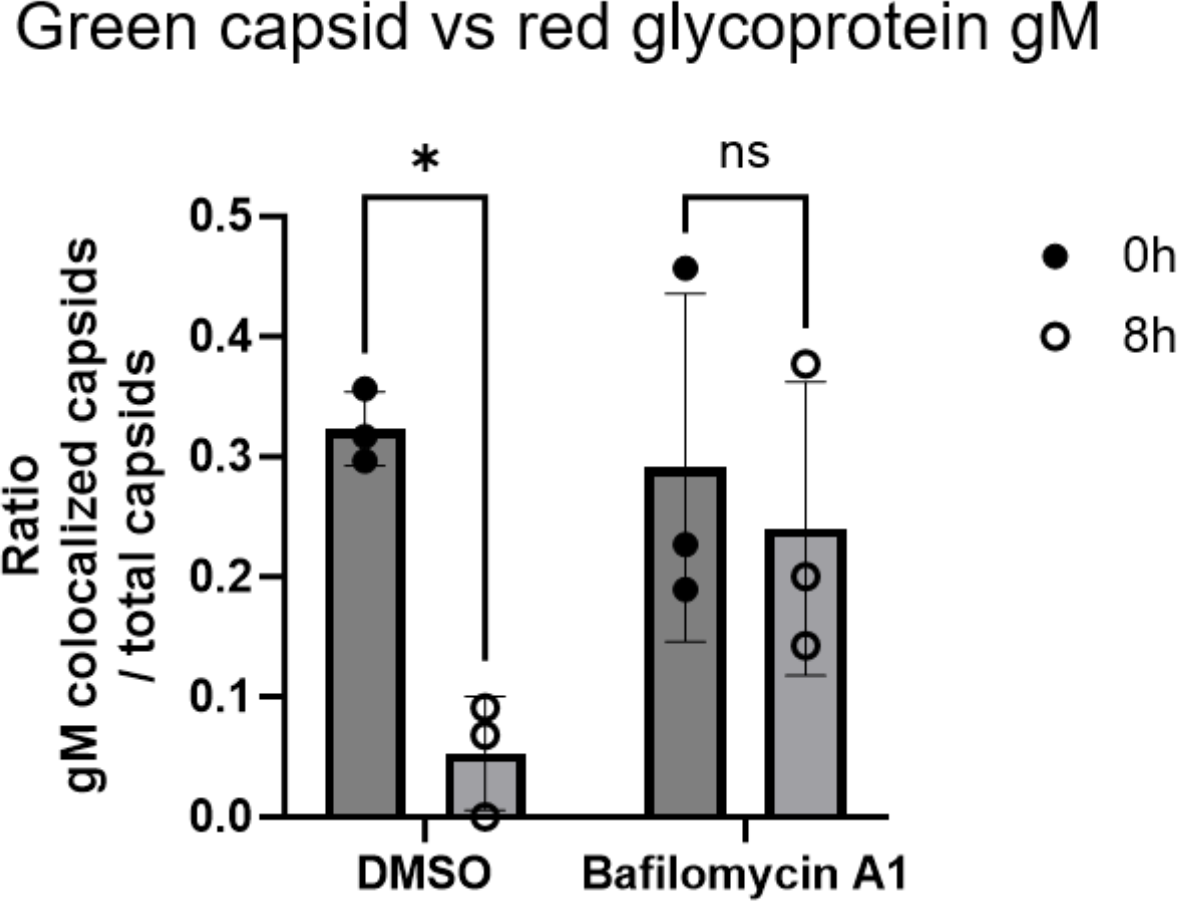
Swapping the position of mNeonGreen and mScarletH has no effect on measured fusion activity and sensitivity to Bafilomycin A1. The capsid was tagged with mNeonGreen and gM with mScarletH, KSHV^mN65-39mS^, (reversed order) and infection was carried out in the presence of solvent control or 50 nM Bafilomycin A1. The ratio of gM-mScarlet colocalized mNeonGreen-ORF65 capsids over total mNeonGreen-ORF65 capsids was calculated. Error bars represent the standard deviation, n=3. ns non-significant; * p<0.05; two-way ANOVA with Sidak’s correction for multiple comparison.

### RRV carrying KSHV gB loses the ability to fuse at the plasma membrane and depends on vesicle acidification for infection

RRV infection is to a large degree sensitive to inhibitors of vesicle acidification [26], but its core fusion machinery is considerably more fusogenic in cell-cell fusion assays than that of KSHV (Figure 1).

Therefore, we wanted to test whether RRV could bypass the requirement of vesicle acidification under conditions of high receptor expression. Indeed, when Raji cells expressed high amounts of the fusion receptor EphB3 after lentiviral transduction, RRV infection occurred independently from vesicle acidification as demonstrated by resistance to Bafilomycin A1 treatment (Fig. 5, filled circles). Next, we wanted to test whether replacing RRV gB with KSHV gB in the virus would change this phenotype, i.e. cause a loss-of-function with regard to virus-cell fusion. As the gH, gL, and gB glycoproteins of KSHV and RRV (Fig. 1) or also EBV [28] function together in cell-cell fusion assays we constructed a hybrid RRV with KSHV gB in place of the original RRV gB, which proved to be replication competent (not shown). Indeed, when we exchanged the RRV gene coding for gB with the gene coding for KSHV gB, the chimeric virus lost the ability to infect independently from vesicle acidification and became susceptible to Bafilomycin A1-mediated inhibition (Fig. 5, open boxes).

**Figure 5.**
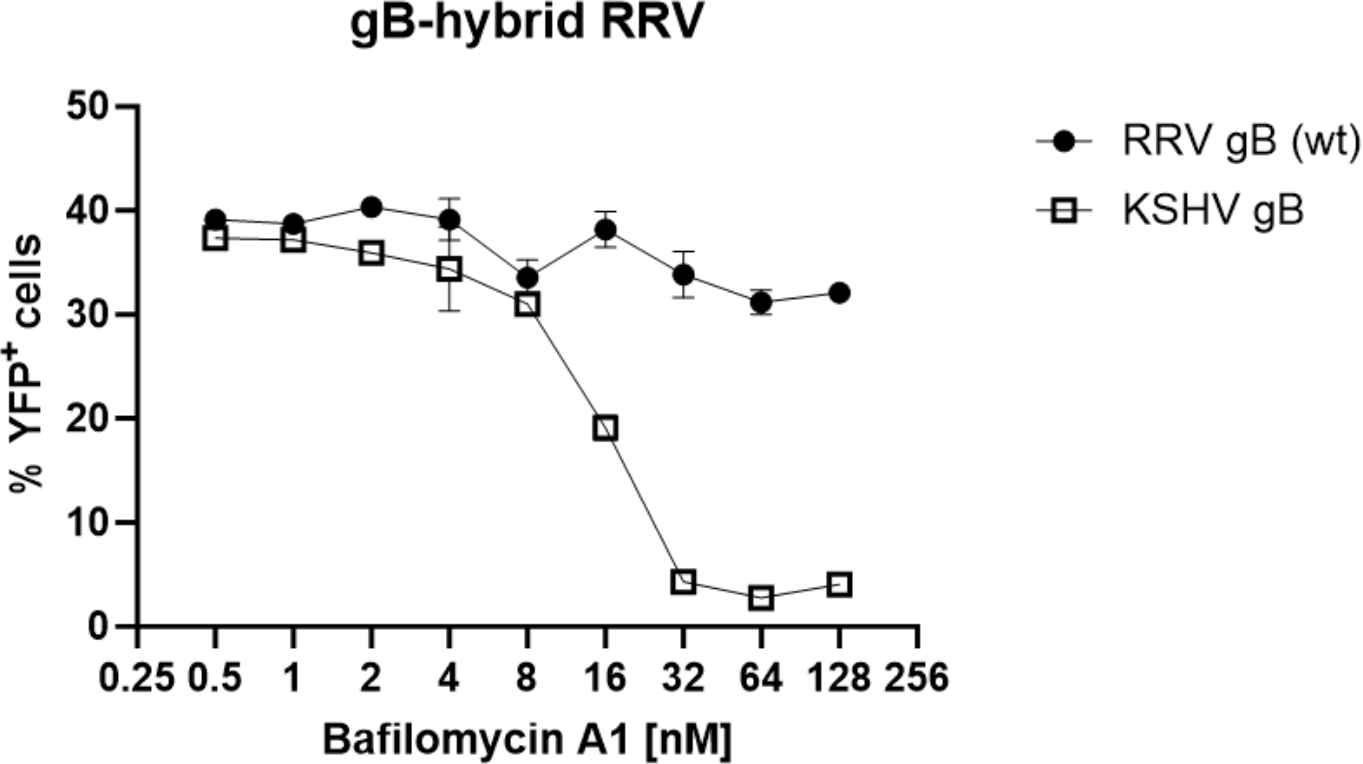
gB dictates sensitivity to inhibition of vesicle acidification. Raji cells that had been transduced to express the RRV gH/gL receptor EphB3 were infected with RRV-YFP with wt gB or with RRV-YFP expressing KSHV gB instead of RRV gB. Infection was quantified as the percentage of YFP^+^ cells.

## DISCUSSION

Using two novel molecular tools, a dual-fluorescent KSHV and a hybrid RRV expressing KSHV gB, we demonstrate that KSHV gB exhibits low fusogenicity and depends on vesicle acidification for the execution of fusion of the viral membrane with cellular membranes. While the potency of inhibitors of vesicle acidification against KSHV infection was well known, we now demonstrate conclusively that the determinant of this sensitivity is the fusion process. Our hybrid RRV expressing KSHV gB further demonstrates that specifically glycoprotein B, the conserved herpesvirus fusion executor, dictates the use of an endocytic entry route, while wt RRV gB allows entry though a pathway that is independent from vesicle acidification when high levels of a gH/gL fusion receptor are present. It should be noted though, that reaching a low pH environment on its own is likely not sufficient for efficient KSHV infection as KSHV gH/gL retains the ability to activate other gB glycoproteins very efficiently upon receptor binding, suggestive of a role of this glycoprotein complex in controlling the fusion process. It is tempting to speculate why KSHV has evolved towards such low fusogenicity. This is even more interesting when one compares KSHV to RRV. While RRV seems to have developed or retained the ability to fuse without the need for vesicle acidification at least in part, KSHV evolved towards an exclusively endocytotic entry pathway. One possible explanation may be minimization of tissue damage and host immune response – a syncytial phenotype might be associated with increased pathogenicity and possibly also with increased genome instability and apoptosis of the host cell [41], a trait that may be selected against when life-long persistence with intermittent or even continuous virus production and shedding is the pathogen’s overarching evolutionary niche.

## METHODS

### Cell culture

Human embryonic kidney (HEK) 293T cells (ATCC) were kindly provided by the laboratory of Stefan Pöhlmann, German Primate Center— Leibniz Institute for Primate Research, Göttingen, Germany. SLK [42] cells (RRID:CVCL_9569) were originally obtained from the (NIH AIDS Research and Reference Reagent program). Rhesus monkey fibroblasts (RF) were kindly provided by the laboratory of Rüdiger Behr, German Primate Center—Leibniz Institute for Primate Research, Göttingen, Germany. If not stated otherwise, all cells were cultured in Dulbecco’s Modified Eagle Medium (DMEM), high glucose, GlutaMAX, 25mM HEPES (Thermo Fisher Scientific) supplemented with 10% fetal bovine serum (FBS) (Serana Europe GmbH), and 50μg/ml gentamycin (PAN Biotech) (D10). iSLK cells[43] were maintained in D10 supplemented with 2.5μg/ml puromycin (InvivoGen) and 250μg/ml G418 (Carl Roth), for selection of iSLK carrying KSHV BACs 200μg/ml hygromycin was added. HUVEC (PromoCell, Heidelberg, Germany), were maintained in standard Endothelial Cell Growth Medium 2 (PromoCell, Heidelberg, Germany). Raji cells (RRID:CVCL_0511) (laboratory of Jens Gruber), were cultured in RPMI (Thermo Fisher Scientific) supplemented with 10% FCS and 50μg/ml gentamycin.

### Lentiviral transduction

For production of lentiviral particles, semi-confluent HEK 293T cells in 10cm cell culture grade petri dishes were transfected with 1.4μg pMD2.G (VSV-G envelope expressing plasmid, a gift from Didier Trono (Addgene plasmid #12259), 3.6μg psPAX2 (Gag-Pol expression construct, a gift from Didier Trono (Addgene plasmid #12260), and 5μg of lentiviral expression constructs using PEI as described before [30]. The supernatant containing the pseudotyped lentiviral particles was harvested 2 to 3 days after transfection and filtered through 0.45μm CA membranes (Millipore). For transduction, lentivirus stocks were used at a 1:5 dilution unless stated otherwise. After 48h, the selection antibiotic blasticidin (Invivogen) was added to a final concentration of 10μg/ml. After initial selection the blasticidin concentration was reduced to 5μg/ml or removed (fusion assay) depending on the experiment.

### Recombinant viruses

Recombinant dual fluorescent viruses were based on BAC16 with the GFP open reading frame replaced with a Zeocin resistance gene and on this BAC carrying a mNeonGreen-GS-ORF65 fusion protein as described earlier, here abbreviaten BAC16 mN65 [35]. Using the technique by Tischer et al. [44], mNeonGreen was replaced with the mScarletH open reading frame to create BAC16 mS65. The mNeonGreen or mScarletH open reading frames were then C-terminally fused to the ORF39 open reading frame, creating BAC16 mS65-39mN and BAC16 mN65-39mS using Primers Bac16_ORF39-GSA-mScarletHmNEON_s ATGAAAGTGACAGTGAAATCGACGAAACGCAAATGATATTCATTGGAAGCGCTGTGAGCAAGGGCGAGG and Bac16_ORF39-GSA-mScarletHmNEONas ATGGAGGAAGAGGGATGGGTTTATAATGCCAATATATCAGCTACTTGTACAGCTCGTCCATGCC to insert the respective fusion protein open reading frame containing the pEPkan-S selection/recombination cassette and a sequence repeat for insertion and scarless excision of the selection marker as described by Tischer et al. [44]. RRV-YFP (MN488839.2)[26,29] was modified using a recombination cassette generated from recombination template AX438 - pcDNA6-KSHVgB-KanS_shuttleplasmid using primers “KSHV gB from aa6 on to RRV” TTTAAAGACCTGTACGCTCTTCTGTACCATCACCTGCAACTGTCCGACGGCCATGATGATAACTAACagattggccaccctgggg and “KSHV gB to RRV rev” GGGTGCGCGAATCGATTGGCCGCGCGGCTCTGGCGGGCGGCAAGTACAGGCGTGTGGGTGTGGTTActcccccgtttccggactg to generate RRV-YFP_KgB, replacing the orf8 gene with that of KSHV. In order to preserve the overlapping RRV ORF7, the first five amino acids of KSHV gB were replaced with those of RRV gB, which according to SignalP prediction does not alter signal peptide functionality [45]. AX438 encodes KSHV gB and the recombination cassette of pEPkan-S [44] after nucleotide 1560, followed by a 87 nucleotide repeat (nucleotides 1478-1560). Sequences of all viruses were confirmed by Illumina-based next generation sequencing.

### Plasmids

#### Infection assays and flow cytometry

The Raji cells in suspension were directly infected with the indicated amount of virus after being seeded in 96-well plates (25,000 cells/well). The suspension cells were centrifuged briefly, the supernatant was discarded and the cells were re-suspended in PBS. Then, the cell suspension was transferred to an equal volume of PBS supplemented with 4% methanol-free formaldehyde (Carl Roth) for fixation. For assessing Bafilomycin A1 inhibitory activity on Raji infection, the cells were plated at 96-well plate, 20,000 cells/well in 50 μL. Then, 50 μL Bafilomycin at twice the indicated concentration was added to cells; the final concentration given in the figure was the one used during incubation. After 30 min incubation, the relevant virus was added to the cells. The virus volume was 25 μL, so the concentration of Bafilomycin A did not change by more than 25% between pre-incubation and post-infection. At 48 h post-infection, cells were harvested and analyzed by flow cytometry to determine the percentage of YFP-positive cells.

For all infection assays a minimum of 5,000 cells were analyzed per well for YFP expression on an ID7000 spectral analyzer (Sony). For immunofluorescence, cells were seeded at approx. 75 000 cells (half that number for HFF) per well on 12-mm coverslips (YX03.1; Carl Roth) in 24-well plates. For experiments using inhibitors, incubation with virus suspension was performed in the presence of inhibitors or solvent controls at the indicated concentrations and medium was exchanged to D10 containing inhibitors or the respective solvent controls. On day two, the plate was cooled down on ice, followed by inoculation with 500 or 1000μl cold virus suspension per well and 30min centrifugation at 4122g and 4°C, followed by another 10min at 4°C. Then, the cells were washed three times in ice-cold PBS, followed either by fixation or medium exchange to D10 and incubation for the indicated timepoints at 37°C in a cell culture incubator, followed by one wash in PBS and fixation. The cells were fixed in 4% methanol-free formaldehyde in PBS for 10 min.

#### Fluorescence microscopy and image analysis

Immunofluorescence was performed as described previously [35]. Z-stacks were imaged using ZEN software (Zeiss) and a laser scanning microscope (Zeiss SLM 800), frame size 2048px x 2048px, scan speed 5, pinhole was set using the 1AU setting provided by the Zeiss software, interval 0.31μm, one separate scan for the mScarletH channel, one for the mNeonGreen channel, and one for the Hoechst channel. For each image, first green positive regions of interest (ROI)s were defined and then red capsids were quantitated using the “find maxima” function of ImageJ in the whole image and in the green (orf39-mNeonGreen)-positive ROIs.

For analysis of figure 3, thresholding was run for mScarletH and mNEONgreen channels using uninfected cells as controls. The Colocalization Plugin (Pierre Bourdoncle) was used for identification of colocalized pixels in each image of the captured stacks. Generated single 8-bit images within the stacks were projected to one image using “Z project, max intensity” and colocalized clusters of pixels representing viral particles were counted using the “Find Maxima” function implemented in Fiji/ ImageJ. Counting of single positive particles was performed using the same function on images after thresholding.

For analysis of figure 4, colocalized regions in maximum projections from three independent stacks were identified as ROI using the “Colocalization Finder” Plugin by Philippe Carl. This technique was chosen as only minimal or no overlap of particles was observed in different z-planes. Limits for the ROIs were set so that no colocalization was observed in controls without virus or for single-fluorescent virus. The ROI containing the colocalized signals was then copied to the original image, and viral particles within this ROI were quantified using the “find maxima” function of ImageJ in the green channel image (in case of mNeonGreen-orf65/orf39-mScarletH particles), the same was also done for the whole image. The ratio of colocalized capsids to total capsids was then calculated for each image and averaged.

Live cells in culture were imaged on a Zeiss AxioVert.A1 cell culture microscope with LED fluorescence imaging.

#### Cell-cell fusion assay

Raji target cells were stably transduced with lentiviral constructs encoding EphB3, or Plxdc2, or EphA2, respectively. After selection, transduced Raji cells were again transduced with a lentiviral construct encoding a Gal4 response element-driven TurboGFP-luciferase reporter for 48 h similar to as described before [35]. 293T effector cells were seeded in 96-well plates at 30,000 cells/well. One day after, 293T effector cells were transfected with a plasmid, encoding the Gal4 DNA binding domain fused to the VP16 trans-activator (VP16-Gal4), and the indicated viral glycoprotein combinations or a pcDNA empty vector control (VP16-Gal4: 31.25 ng/well, RRV gH: 12.5 ng/well, RRV gL: 62.5 ng/well, RRV gB: 18.75 ng/well, KSHV gH: 12.5 ng/well; KSHV gL: 62.5 ng/well, KSHV gB: 18.75 ng/well, pcDNA only: 93.75 ng/well) using PEI as described before [46]. 24 h after transfection, the medium with the transfection mix was removed and replaced with 100 μL fresh D10. The target cells were counted, and 40,000 target cells were added to 293T effector cells in 100 μL fresh R10. Triplicate wells were analyzed for all target-effector combinations. After 48 h, cells were washed once in PBS and lysed in 35 μL 1× Luciferase Cell culture lysis buffer (E1531, Promega) for 15 min at room temperature and centrifuged for 10 min at 4°C. A volume of 20 μL of each cell lysate was used to measure luciferase activity using the Beetle-Juice Luciferase Assay (PJK), according to the manufacturer’s instructions on a Biotek Synergy 2 plate reader.

## ACKNOWLEDGEMENTS

We thank Stefan Pöhlmann for support, Marija Backovic and Christian Münz for helpful discussions, Rüdiger Behr for rhesus monkey fibroblasts, Klaus Korn for human foreskin fibroblasts.

This work was funded by grants to ASH from the Deutsche Forschungsgemeinschaft (grants HA 6013/4-1, DFG HA 6013/10-1), by a grant from the Wilhelm-Sander Foundation (https://www.wilhelm-sander-stiftung.de; project no. 2019.027.1) to ASH. S.L. was funded by a scholarship of the China Scholarship Council (CSC), file number 202106300006.

